# Repurposing statin and L-glutamine: replenishing β-cells in hyperlipidemic T2D mouse model

**DOI:** 10.1101/2021.04.22.440866

**Authors:** Sayantani Pramanik Palit, Roma Patel, Nishant Parmar, Nirali Rathwa, Nilay Dalvi, A. V. Ramachandran, Rasheedunnisa Begum

**Author notes:** Corresponding author, Department of Biochemistry, Faculty of Science, The Maharaja Sayajirao University of Baroda, Vadodara, Gujarat, India.

## Abstract

While cases of obesity-induced type 2 diabetes (T2D) are on the upswing, current therapies help only manage the symptoms. Of late, L-glutamine has been implicated in the amelioration of T2D by virtue of its glucagon like peptide-1 (GLP-1) secretagogue property. Alongside, there are mixed reports on adiponectin (insulin sensitizer) potentiating property of statins. We aimed to investigate the effect of pitavastatin (P) and L-glutamine (LG) combination on glycemic control and pancreatic β-cell regeneration in a high-fat diet (HFD)+Streptozotocin (STZ) induced T2D mouse model. C57BL6/J mice treated with HFD+STZ were randomly assigned into four groups: Diabetic control, L-glutamine, Pitavastatin and P+LG. Control group was fed with the chow diet. Significant amelioration in insulin resistance along with plasma glucose, lipid profile, adiponectin levels, and mitochondrial complexes I, II, III activities were observed in P+LG group as compared to HFD+STZ treated group. Phosphoenolpyruvate carboxykinase, glucose 6-phophatase, glycogen phosphorylase, and GLUT2 transcript levels were reduced with increased glycogen synthase transcript levels in liver. Further, the protein levels of Insulin receptor 1-β, pAkt/Akt, and AdipoR1 were restored in the skeletal muscle and a significant increase in islet number as a result of β-cell regeneration and reduced β-cell death were also observed in the combination drug treated group. Thus, L-glutamine and pitavastatin in combination can induce β-cell regeneration and regulate glucose homeostasis to bring about amelioration of HFD+STZ induced T2D.

## 1. Introduction

Insulin resistance, reduced insulin secretion, increased hepatic glucose output, dyslipidaemia/ hypercholesterolemia, and β-cell degeneration create the landscape of type 2 diabetes (T2D) (Pramanik et al., 2017). Many therapeutic modalities have been investigated to manage hyperglycaemia and, the choice of drug varies from case to case. While the initial borderline or pre-diabetic state management is relatively easy by incorporating lifestyle and dietary interventions to boost insulin sensitivity, a progressed state of developed T2D needs to be addressed from a β-cell regenerative view point. Most often at diagnosis, islet function may already be reduced by 50% and β-cell mass by 60%. The reduction in β-cell mass is due to accelerated apoptosis (Garber, 2011). While, autopsy studies conducted on T2D cadavers have shown 0-65% loss in β-cells (Matveyenko and Butler, 2008), Butler et al. showed that obese patients in pre-diabetic and T2D state had 40% reduction in β-cell mass when compared to obese individuals without T2D in cadaver pancreatic autopsy (Butler et al., 2003). Only a handful of current treatment modalities address the worsening condition of β-cell loss with time. However, we believe that the currently available on-shelf drugs have untapped potentials to reverse and restore glucose-homeostasis. Enhancing the incretin hormone GLP-1 level is one such intervention. It has a number of protective effects on the β-cells, including a reduction in apoptosis and enhancement of β-cell proliferation and neogenesis (Garber, 2011). L-glutamine is known to stimulate glucagon-like peptide 1 (GLP-1) secretion in human subjects by raising Cytosolic Ca^2+^ and cAMP in intestinal L-cells (Tolhurst et al., 2011). Interestingly, while in prediabetics L-glutamine levels have been reported to be higher (Owei et al., 2019), it has been found to be low in T2D subjects (Tsai et al., 2012; Chen et al., 2019). Another class of drugs, statins notoriously known to induce new onset diabetes but are used for the comprehensive management of hypercholesterolemia associated with obesity induced T2D (Kim et al., 2019). Pitavastatin has shown a remarkable outcome on the glycemic control in patients with T2D compared to other members of the statin group (Mita et al., 2013, Zhao and Zhao, 2015).

Thus, going ahead we proved our thesis that the additive/synergistic effect resulting from combining two small molecules together i.e., pitavastatin with a GLP-1 secretagouge, L-glutamine to ameliorate high fat diet (HFD) and streptozotocin (STZ) induced T2D in C57BL6/J mice can ameliorate T2D manifestations.

## 2. Materials and Methods

### 2.1 Animals and experimental strategy

9-10 weeks old (22 grams) male C57BL/6 mice procured from Advanced Centre for Treatment, Research and Education in Cancer (Navi-Mumbai, India) were housed in our animal house (Dept. of Biochemistry, The Maharaja Sayajirao University of Baroda, Vadodara, Gujarat, India), and maintained in a 12-h light-dark cycle, with free access to standard chow/HFD (Keval Sales Corporation, India) and water supplied through a water bottle.

Based on the plasma levels of glucose and body weight, the animals were divided first into two balanced groups; 1) fed a standard chow diet (n=6) and another HFD (n=25) to develop obesity induced insulin resistance. After 20 weeks the HFD fed mice were administered three consecutive low doses of streptozotocin (STZ) (MP Biomedicals, India) (40 mg/kg i.p.) to induce β-cells loss (Parilla et al., 2018). HFD+STZ induced T2D was confirmed with body weight (BW) ≥30 grams and three consecutive readings of fasting blood glucose (FBG) ≥240 mg/dL (Fig S1A).

After 22^nd^ week the HFD+STZ group was further divided into four groups (n=6/group) namely 2) HFD/STZ, 3) Pitavastatin (P, 0.5mg/kg b.w in diet) (Zambarakji et al., 2006), 4) L-glutamine (LG, 500mg/kg b.w p.o) (Badole et al., 2014; Sadar et al., 2016) and 5) Pitavastatin (0.5mg/kg b.w in diet)+ L-glutamine (P+LG, 500mg/kg b.w p.o) treated (Fig S1B). The treatment was continued daily for 6 weeks along with 5-bromo-2’-deoxyuridine (BrdU) (MP Biomedicals, India) on alternative days (100mg/kg bw i.p.). The FBG and BW of the animals were measured weekly by glucometer (TRUEresult® - Nipro) and weighing scale respectively along with the food and water intake. At the end of 6 weeks of treatment, glucose and insulin tolerance were evaluated by Intraperitoneal glucose tolerance test (IPGTT) and Intraperitoneal insulin tolerance test (IPITT). Mice were fasted for 6h and injected with glucose (2g/kg b.w) and insulin (0.5U/kg) for IPGTT and IPITT procedure (Jørgensen et al., 2017; Guo et al., 2018). A gap of two days was given after completing glucose tolerance and insulin sensitivity test when the drug treatments were continued. Animals were put on fasting for 6 hours and 800-900µL blood was collected by orbital sinus and animals were sacrificed for tissue collection. Plasma was separated and was stored for further biochemical parameter evaluation (Fig S1A).

All the procedures followed the institutional guidelines approved by Institutional Ethical Committee for Animal Research (IECHR), Faculty of Science, The Maharaja Sayajirao University of Baroda, Vadodara, Gujarat, India (MSU/BIOCHEMISTRY/IAEC/2018-21).

### 2.2 Metabolic and Biochemical Assessment

#### 2.2.1 Lipid Profiling

Plasma separated was used for lipid profiling (total cholesterol, triglyceride, high density lipoprotein) as per our previous study (Palit et al., 2020). Friedewald’s (1972) formula was used for calculating low density lipoprotein (LDL) (Knopfholz et al., 2014).

#### 2.2.2 Assessment of plasma insulin and adiponectin

The plasma levels of insulin and adiponectin of the experimental animals were determined by the mouse ELISA kit (RayBio, USA) as per manufacturer’s instructions.

### 2.3 Assessment of transcript levels

Liver and skeletal muscle were collected and stored in RNAlater™ Stabilization Solution (Thermo Fisher Scientific, USA) and total RNA was extracted by the Trizol method as described previously (Pali et al., 2020). The expression of targeted genes and *GAPDH* transcripts were monitored by LightCycler®480 Real-time PCR (Roche Diagnostics GmbH, Germany) using gene-specific primers (Eurofins, India) as shown in Table S1. Expression of *GAPDH* gene was used as a reference. Real-time PCR was performed as described previously (Palit et al., 2020).

### 2.4 Mitochondria isolation from skeletal muscle and estimation of oxygen consumption rate (OCR)

Mitochondria was isolated from skeletal muscle using mitochondria isolation kit (Thermo Scientific ™, Catalog no. 89801) using manufacturer’s protocol. The isolated mitochondria were resuspended in mitochondria respiration buffer (110mM Sucrose, 0.5mM EGTA, 70 mM KCl, 0.1% FFA free BSA, 20 mM HEPES, 3 mM MgCl_2_, and 10 mM KH_2_PO_4,_ 20mM Taurine) (Butterick et al, 2016). Mitochondrial outer membrane integrity of the isolated mitochondria was assessed by impermeability to exogenous Cytochrome c which was consistently greater than 95%. Respiratory chain complexes I-III activities were recorded using 100 mM Pyruvate & 800 mM Malate (complex I), 1M Succinate (complex II) and 10 mM α-glycerophosphate (complex III) as the respective substrates (Li and Graham, 2012) and protein concentration was estimated by Bradford method (Jones et al., 1989). All chemicals were purchased from Sigma-Aldrich, USA. OCR was determined by measuring the amount of oxygen (nmol) consumed, divided by the time elapsed (min) and the amount of protein present in the assay (Li and Graham, 2012).

### 2.5 Pancreatic tissue preparation, immunohistochemistry (IHC), assessment of β-cells regeneration and apoptosis

At the end of the experiment, animals were sacrificed and pancreatic tissues were fixed in 10% formalin and was be processed for paraffin embedding. 5µm sections were prepared from the blocks. These sections were deparaffinised with 100% xylene and rehydrated by ethanol gradient wash (100%, 95%, 80% and 70%) and proceeded with an antigen retrieval step (1N HCl at 37°C for 45 min). Subsequently, these sections were blocked with 5% donkey serum (Jackson Immuno Research Laboratories, Inc. USA). Antibodies were diluted in the blocking solution as indicated in table S2. The sections were incubated with primary antibody for 1h at 37°C, washed with PBS and incubated with secondary antibody for 45 min at 37°C. These sections were also stained with DAPI with anti-fade mounting medium, DAPI Gold Antifade (Thermo Scientific). The slides were visualized under confocal microscope (Olympus FV10i, Olympus, Tokyo, Japan) at 60X and images were processed using Fluoview. Co-staining (immunofluorescence) with anti-insulin and anti-BrdU was considered for proliferation of β-cells. Co-staining with anti-insulin, anti-PAX-4 (Paired box 4), and anti-ARX (Aristaless-related homeobox-encoding gene) was considered for transdifferentiation while anti-insulin, anti-NGN-3 (Neurogenin 3) and anti-PDX-1 (Pancreatic and Duodenal Homeobox 1) was considered for neogenesis. β-cells death was evaluated by the co-staining of insulin and Terminal deoxynucleotidyl transferase dUTP nick end labelling (TUNEL) as well as insulin and Apoptosis Inducing Factor (AIF) markers. Results are expressed as the percentage of specified markers for regeneration and death.

### 2.6 Western blot analysis

Skeletal muscle was collected post sacrifice and stored in -80°C. The tissue was homogenized in liquid nitrogen and lamelli buffer containing urea (1M Tris HCl, 10%SDS, 20% glycerol and 10% β-mercaptoethanol. To this, 1M Urea was added in 1:1 ratio) using mortar pestle. The homogenate was collected and sonicated twice in chilled condition and centrifuged to remove tissue/cell debris. The protein concentration in the lysates were estimated using Bradford method and 25µg of the lysate was resolved on 10% SDS-PAGE, followed by electrophoretic transfer to PVDF membrane. Then immunoblot analysis was processed with targeted primary antibodies and with the secondary antibodies conjugated with HRP (Table S3). The membrane was visualized with the clarity western ECL substrate (Bio-Rad Laboratories, USA) in the ChemiDoc™ Touch Imaging System and blots were analysed in image lab software (Bio-Rad Laboratories, USA).

### 2.7 Statistical analyses

The data analysed are expressed as the mean ± SEM and *p*<0.05 was considered as significant. The difference between groups were conducted by one-way ANOVA followed by the Tukey’s test for multiple group analysis. All the analyses were carried out in GraphPad Prism 5 software.

## 3. Results

### 3.1 Animals and experimental strategy

After 20 weeks of HFD treatment, animals turned obese and insulin resistant. After STZ administration, the FBG levels of these animals surpassed 240 mg/dl (Figure S2A). There was a significant increase in the FBG levels and BW post STZ treatment (Table S2B & Fig. S2C).

### 3.2 Metabolic and Biochemical Assessment

The results showed a significant decrease in the body weight (*p*<0.05) and a significantly decreased FBG (*p*<0.001) in P+LG group at the end of the treatment as compared to HFD+STZ (Fig. 1A and B).

**Figure 1.**
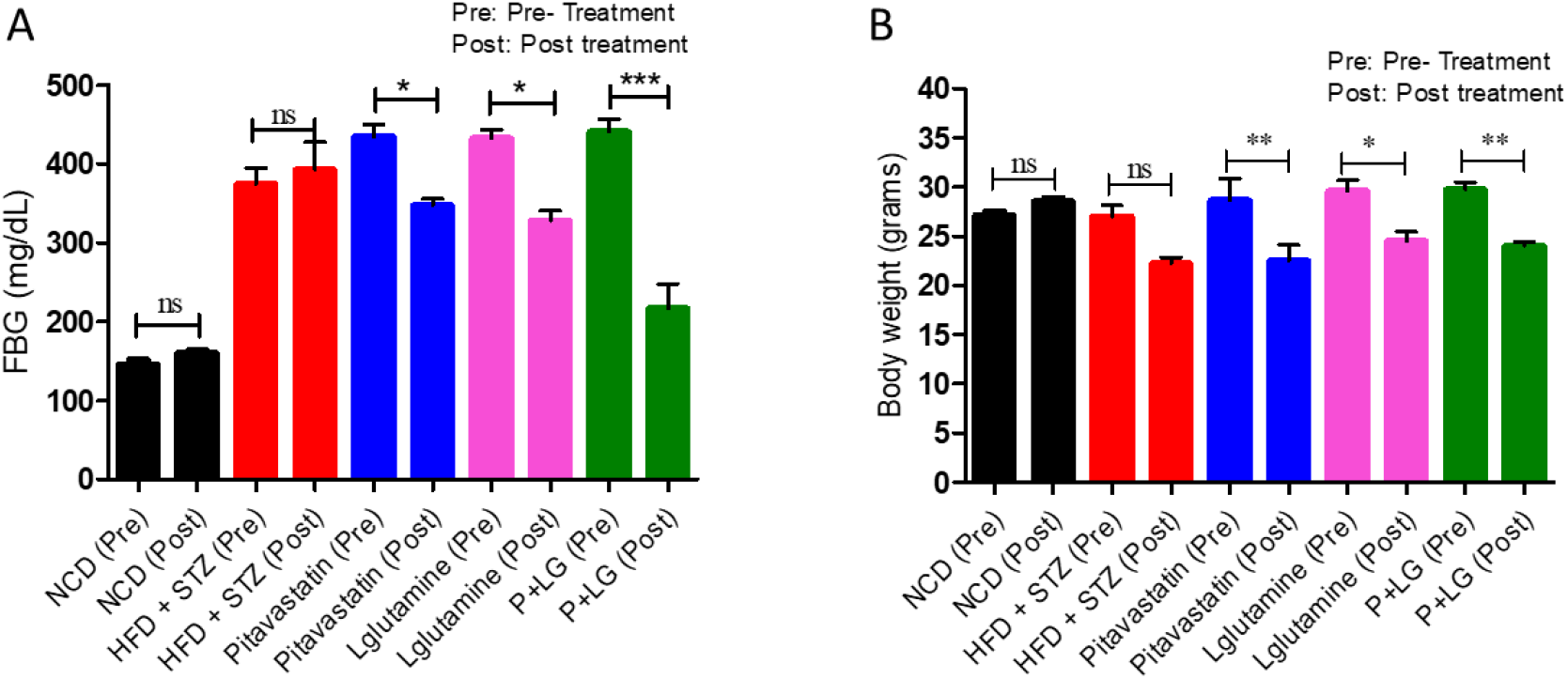
Body weight (BW) and fasting blood glucose (FBG) in treated groups: **A)** A significant decrease in the FBG levels were observed in the monotherapy treated group which was further reduced in the combination P+LG treated group as compared to pre-treatment, **B)** A significant change in body weight was observed in the monotherapy treated and combination treated groups as compared to pre-treatment (ns=non-significant, \**p*< 0.05, \*\**p*< 0.01, \*\*\**p*=0.001, Pre vs. Post; n=5/group).

#### 3.2.1 Intraperitoneal glucose tolerance test (IPGTT) and Intraperitoneal insulin tolerance test (IPITT)

The HFD+STZ and monotherapy treated groups had higher base line glucose after four hours fasting. Blood glucose levels in mice treated with P+LG were significantly lower than in mouse treated with HFD+STZ treated mice at 60, 90 and 120 minutes of glucose administration (Fig.2A). Area under the curve (AUC) 0–120 curve indicated improved glucose tolerance in the P+LG treated groups as compared to HFD+STZ group (Fig.2B). Blood glucose levels in mice treated with P+LG was significantly lower than in mice treated with P, LG and HFD+STZ groups at 60, 90 and 120 minutes of insulin administration (Fig.2C). AUC0–120 curve in mice treated P+LG combination was significantly lower than in HFD+STZ group (Fig.2D).

**Figure 2.**
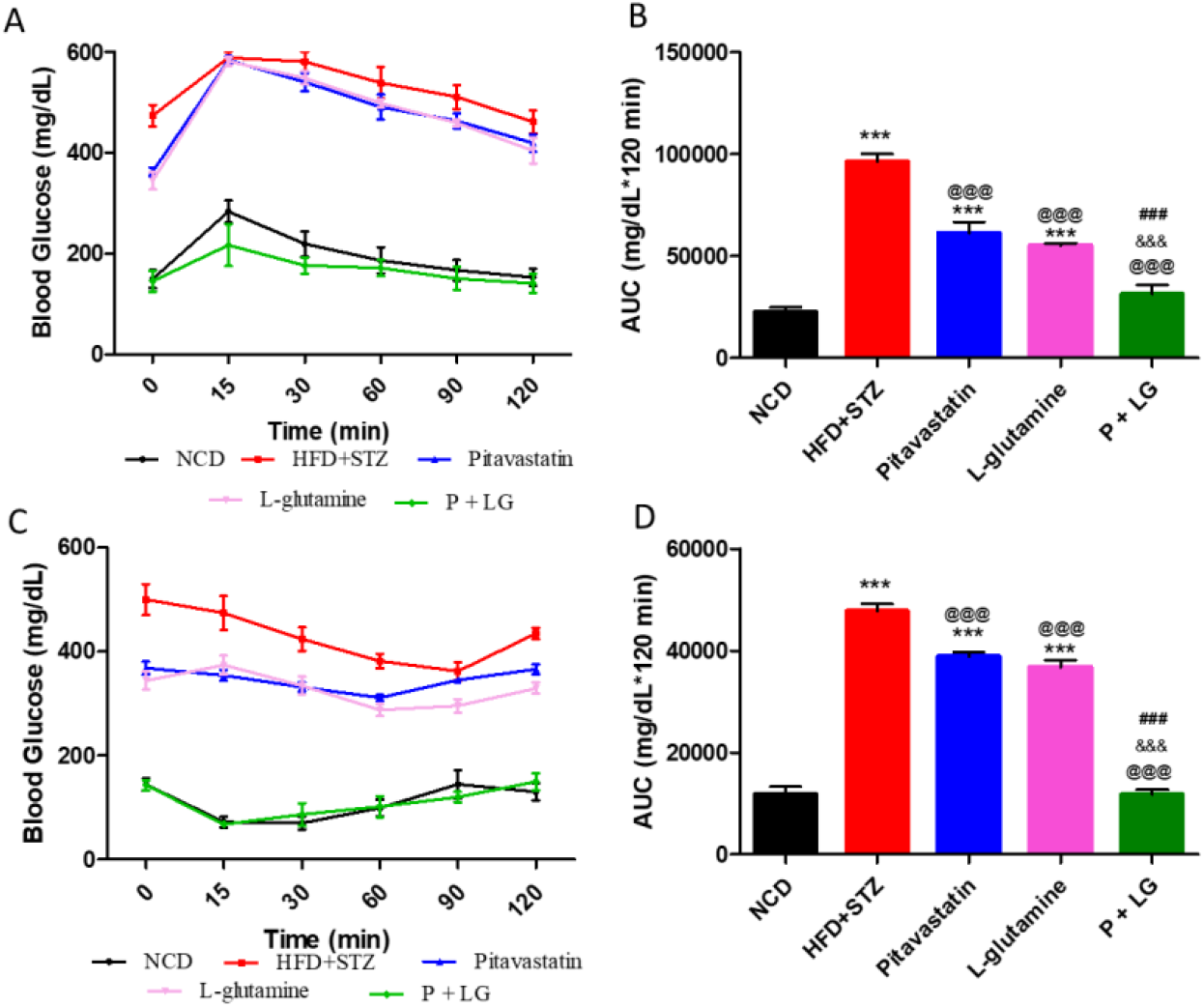
Intraperitoneal glucose tolerance test (A-B) & Intraperitoneal insulin tolerance test (C-D) in the treated animals. Treatment with P+LG improves glucose and insulin tolerance in animals on high-fat diet treated with three low doses of STZ. Panel A shows blood glucose profiles during intraperitoneal glucose tolerance test (IPGTT); Panel C shows blood glucose profiles during intraperitoneal insulin tolerance test (IPTT). Panels B and D show respective areas under blood glucose concentration curve (AUC) from OGTT and IPTT tests (^@@@^ *p*<0.001 vs. HFD+STZ, ^&&&^ *p*<0.001 vs. P, ^###^ *p*<0.001 vs. LG, \*\*\**p*<0.001 vs. NCD, n=5/ group).

#### 3.2.2 Lipid Profiling

Plasma lipid profile indicated improved profile: Triglyceride (TG), Total Cholesterol (TC) and Low density lipoprotein (LDL) levels were significantly reduced in the P+LG treated group as compared to HFD+STZ as shown in Fig. 3A, B & C. High density lipoprotein level was significantly increased in HFD+STZ treated group which was significantly reduced upon combination treatment. (Fig. 3D).

**Figure 3.**
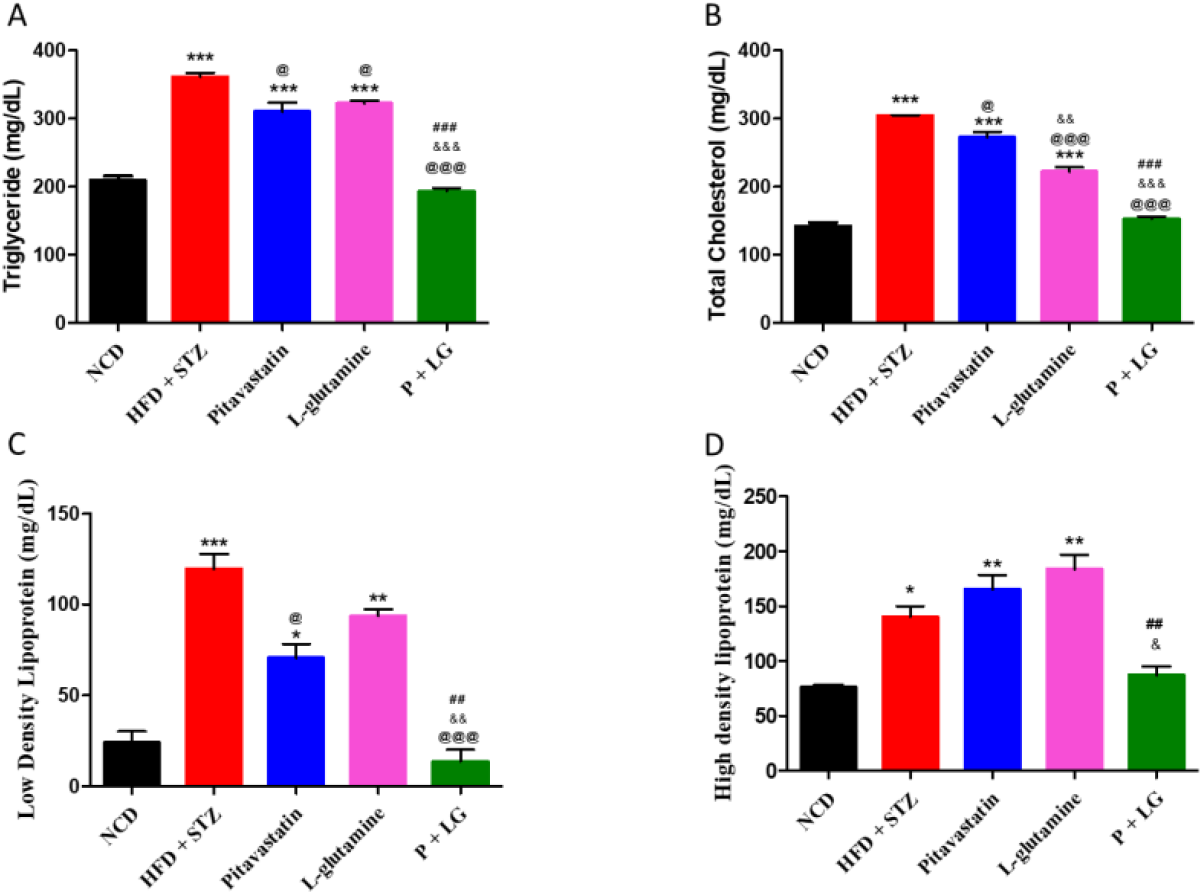
Plasma lipid levels. **A)** Triglyceride level was significantly increased in the HFD+STZ group and was significantly reduced in pitavastatin, L-glutamine and P+LG group, **B)** Total cholesterol level was significantly increased in the HFD+STZ group and was significantly reduced in pitavastatin, L-glutamine and P+LG group, **C)** Low density lipoprotein level was significantly increased in the HFD+STZ group and was significantly reduced only in pitavastatin and P+LG group as compared to HFD+STZ group, and **D)** High density lipoprotein level was significantly increased in HFD+STZ treated group and it was significantly reduced in the combination treated group. (^@^ *p*<0.05, ^@@@^ *p*<0.001 vs. HFD+STZ, ^&^*p*<0.05, ^&&^*p*<0.01, ^&&&^*p*<0.001, vs. LG, ^##^*p*<0.01, ^###^*p*<0.001, vs. LG, \**p*<0.05, \*\**p*<0.01, \*\*\**p*<0.001, vs. NCD, n=5/group).

#### 3.2.3 Plasma insulin and adiponectin levels

There was a significant decrease in insulin and adiponectin levels in the HFD+STZ treated group. However, upon treatment there was a significant increase in the insulin level in the combination treated group (Fig 4 A). There was a significant increase in the adiponectin level in the monotherapies while a normo-adiponectinemea was seen in the P+LG group (Fig 4B).

**Figure 4.**
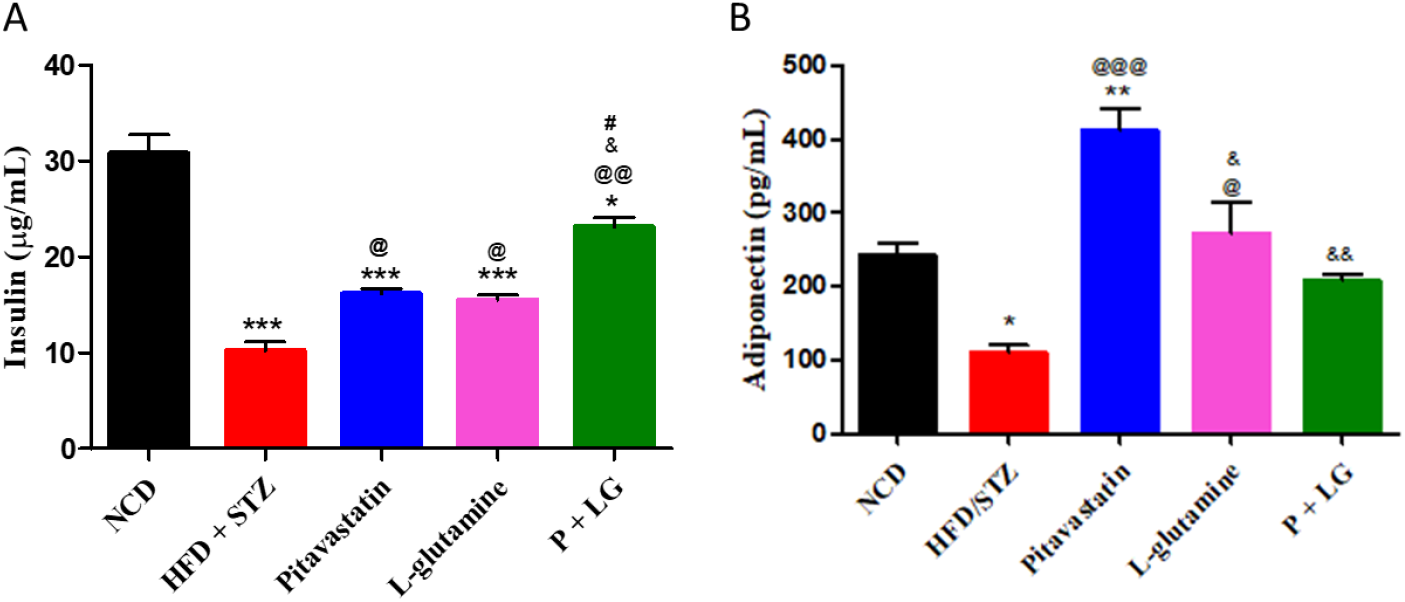
Plasma insulin & adiponectin levels: **A)** Increased insulin was observed in P+LG treated group. **B)** A significantly increased adiponectin was observed in the monotherapy treated groups as compared to HFD+STZ group and normo-adiponectinemia was observed in the combination treated group (^@^*p*<0.05, ^@@^*p*<0.01, ^@@@^*p*<0.001, vs. HFD+STZ, ^&^*p*<0.05, ^&&^*p*<0.01, vs. LG, ^#^*p*<0.05, vs. LG, \**p*<0.05, \*\**p*<0.01, \*\*\**p*<0.001, vs. NCD, n=5/ group).

### 3.3 GLUT2, and glucoregulatory enzyme transcript levels and their activities in liver

The transcript levels and activities of key enzymes involved in the glucoregulatory metabolic pathways were assessed in liver. The results indicate a significantly increase in the fold change and activity of glucokinase [*GCK*] (glycolysis) (*p*<0.001, *p*<0.01; Fig 5A & 5B), increased fold change of g*lycogen synthase* (glycogenesis) (*p*<0.01; Fig 5H), and increased glycogen content (*p*<0.001; Fig 5I) in the combination treated group. Further, there was a significant decrease in the fold change and activities of FBPase and PEPCK (gluconeogenesis) (*p*<0.05, *p*<0.001, *p*<0.001, *p*<0.001 respectively; Fig 5C, D, E & F) along with a concomitant decrease in *G6Pase* fold change (*p*<0.01; Fig 5G). Additionally, the g*lycogen phosphorylase* (glycogenolysis) transcript level fold change were also reduced (*p*<0.001; Fig 5J) in the combination treated group with a concomitant decrease in *GLUT2* receptor transcript level fold change (*p*<0.001; Fig 5K).

**Figure 5.**
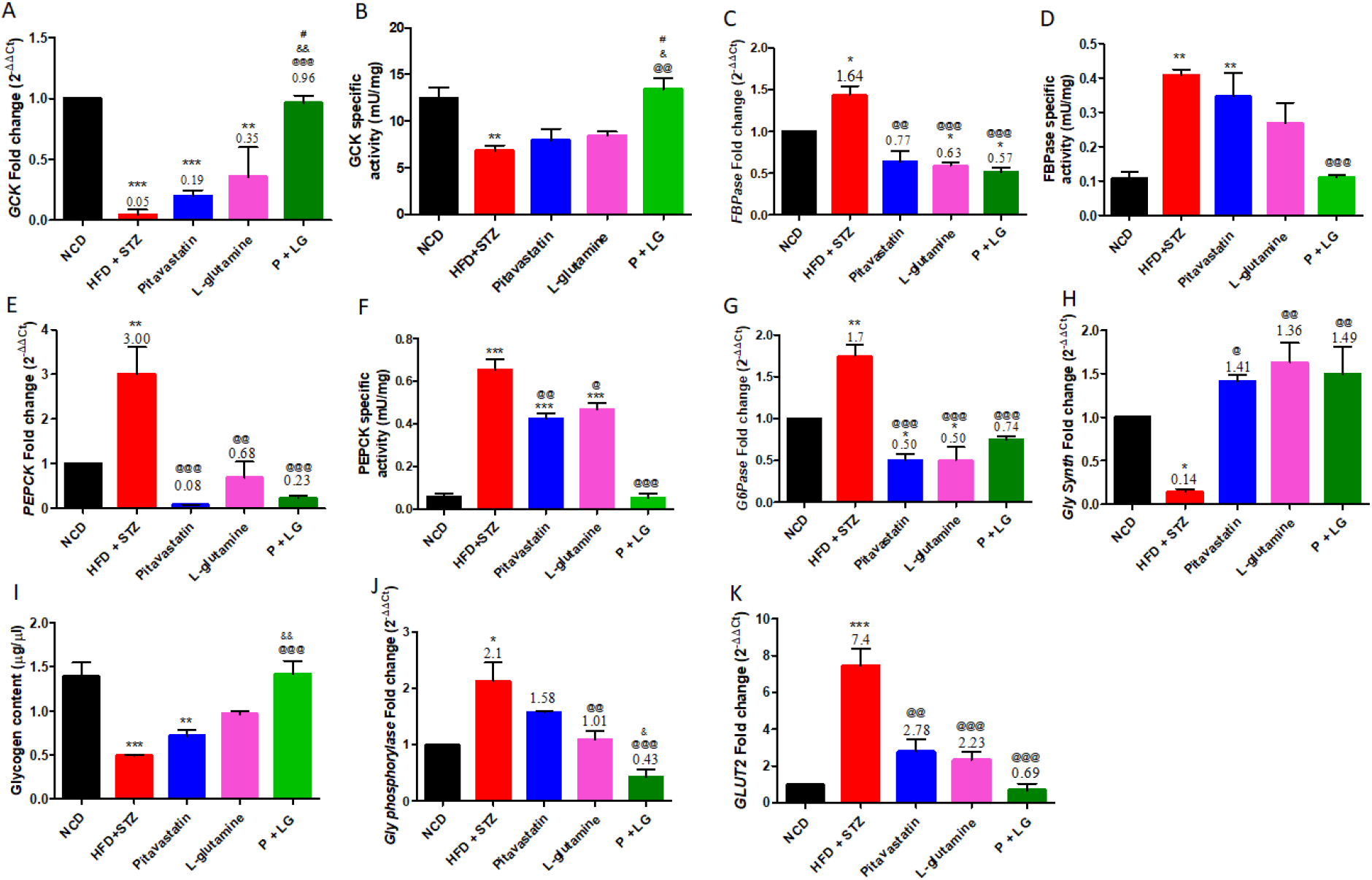
Glucoregulatory enzyme transcript levels in liver: **A) *GCK* (glycolysis) transcript levels:** There was a 0.05 fold decrease in the *GCK* fold change of HFD+STZ group as compared to NCD which was significantly increased in the P+LG treated group, **B) GCK activity:** There was a significant decrease in the activity of GCK in the HFD+STZ treated group. Upon P+LG treatment there was a significant increase in the activity of GCK; **C) *FBPase* (gluconeogenesis) transcript levels:** There was a 1.64 fold rise in the *FBPase* expression of the HFD+STZ group as compared to NCD. The transcript levels were significantly reduced in pitavastatin (0.77 fold), L-glutamine (0.63 fold) and the P+LG (0.57 fold) groups, **D) FBPase activity:** There was a significant rise in the FBPase activity of the HFD+STZ group as compared to NCD. The enzyme activity levels were significantly reduced in the P+LG group as compared to HFD +STZ. **E) *PEPCK* (gluconeogenesis) transcript levels:** There was a 3 fold rise in the *PEPCK* expression of HFD+STZ group as compared to NCD. A significant decrease in the fold change of *PEPCK* transcript levels was observed in pitavastatin, L-glutamine and P+LG treated groups as compared to HFD+STZ, **F) PEPCK enzyme activity**: There was a significant rise in the PEPCK activity of HFD+STZ group as compared to NCD which was reversed upon combination treatment; **G) *G6Pase* (glucose 6 phosphatase) transcript levels:** There was a significant increase in the *G6Pase* transcript levels in the HFD+STZ group as compared to NCD and was significantly reduced upon mono-and P+LG treated groups as compared to HFD +STZ. **H) *Gly Synth* (Glycogenesis) transcript levels:** There was a significant decrease in the expression of *Gly Synth* transcript levels in HFD +STZ treated group as compared to NCD and a significant rise in the *Gly Synth* transcript levels in mono- and P+LG treated group as compared to HFD +STZ; **I) Glycogen content:** The glycogen content was significantly reduced in the HFD+STZ group as compared to NCD. There was a significant increase in the glycogen content in the P+LG treated group as compared to HFD +STZ treated group; **J) *Gly phosphorylase* (glycogenolysis) transcript levels:** There was a significant increase in the *Gly phosphorylase* transcript levels in the HFD + STZ treated group as compared to NCD. The *Gly phosphorylase* transcript levels were significantly decrease in the pitavastatin and P+LG treated groups as compared to HFD +STZ; **K) *GLUT 2* gene transcript levels:** There was a significant rise in the *GLUT 2* expression of HFD +STZ treated group as compared to NCD. There was a significant decrease in the *GLUT 2* transcript levels in pitavastatin, L-glutamine and P+LG treated groups as compared to HFD +STZ. (^@^*p*<0.05, ^@@^*p*<0.01, ^@@@^*p*<0.001, vs. HFD+STZ, ^&&^*p*<0.01, ^&&&^*p*<0.001, vs. LG, ^#^*p*<0.05, vs. LG, \**p*<0.05, \*\**p*<0.01, \*\*\**p*<0.001, vs. NCD, n=4-5/group)

However, the monotherapies; pitavastatin and L-glutamine individually could only decrease gluconeogenesis [*PEPCK & G6Pase*] (*p*<0.001 & *p*<0.01 and *p*<0.01 & *p*<0.05 respectively; Fig 5E & G). The action of monotherapies also reflected in the *GLUT2* receptor by decreasing its transcript level fold change (*p*<0.001; Fig 5K).

### 3.4. Mitochondrial biogenesis marker transcript levels in skeletal muscle

A significant increase in the fold expression of *SIRT 1* was observed in the monotherapies and P+LG treated groups (Fig 6A) along with a significant increase in the fold expression of *PGC 1α* and *TFAM* in the monotherapies and P+LG treated groups (Fig 6B & C).

**Figure 6:**
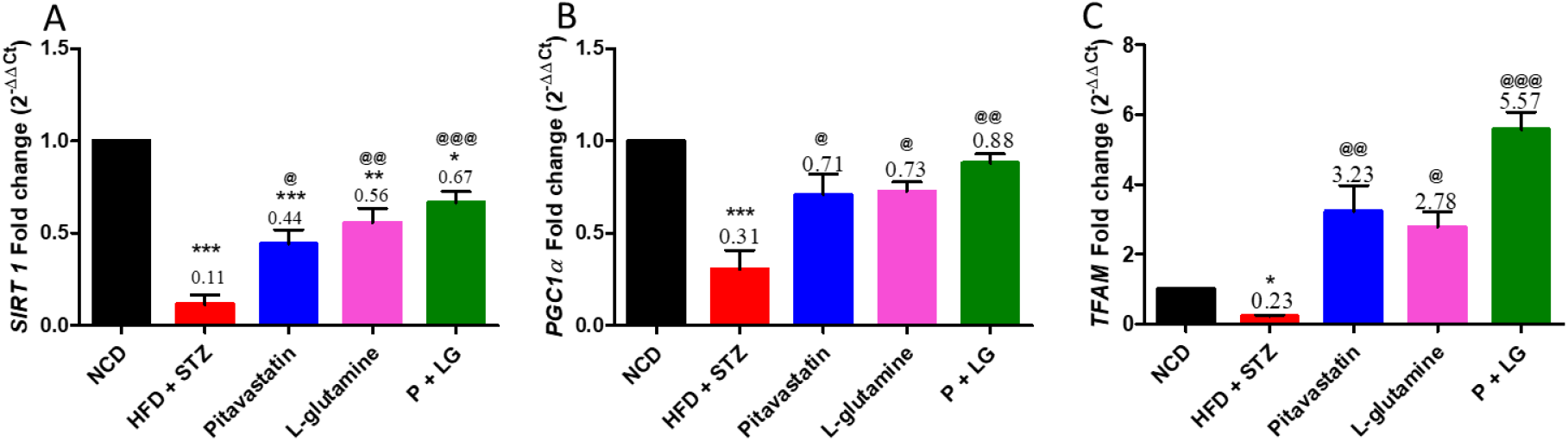
Mitochondrial biogenesis marker transcript levels in skeletal muscle: **A) *SIRT1:*** There was a 0.11 fold decrease in the *SIRT1* transcript levels of HFD+STZ treated group. A 0.44, 0.56 and 0.67 fold increase in the *SIRT1* transcript levels were observed in the pitavastatin, L-glutamine and P+LG treated groups respectively; **B) *PGC1α:*** There was a 0.31 fold decrease in the HFD+STZ treated group significant increase in the transcript levels of *PGC1 α*. The pitavastatin, L-glutamine and P+LG treated groups showed 0.71, 0.73 and 0.88 fold increase in the *SIRT1* transcript levels respectively, and **C) *TFAM*:** There was a 0.23 fold drop in the *TFAM* transcript levels in the HFD+STZ treated group which increased by 3.23, 2.78 and 5.57 fold in the pitavastatin, L-glutamine and P+LG treated groups respectively. (^@^*p*<0.05, ^@@^*p*<0.01, ^@@@^*p*<0.001, vs. HFD+STZ, \**p*<0.05, \*\**p*<0.01, \*\*\**p*<0.001, vs. NCD, n=4-5/group)

### 3.5. Estimation of oxygen consumption rate (OCR)

The OCR of state 3/state 4 in HFD+STZ group was reduced as compared to NCD for the mitochondrial complexes I, II and III. A significant increase in the OCR of pitavastatin, L-glutamine and P+LG groups were observed in CI (^@@^*p*<0.01, ^@^*p*<0.05 & ^@@@^*p*<0.001) (Fig 7A) and CII (^@^*p*<0.05, ^@^*p*<0.05 & ^@@@^*p*<0.001) (Fig 7 B), respectively. However, the OCR of state 3/state 4 in complex III was significantly improved only in the P+LG treated group (Fig 7 C).

**Figure 7:**
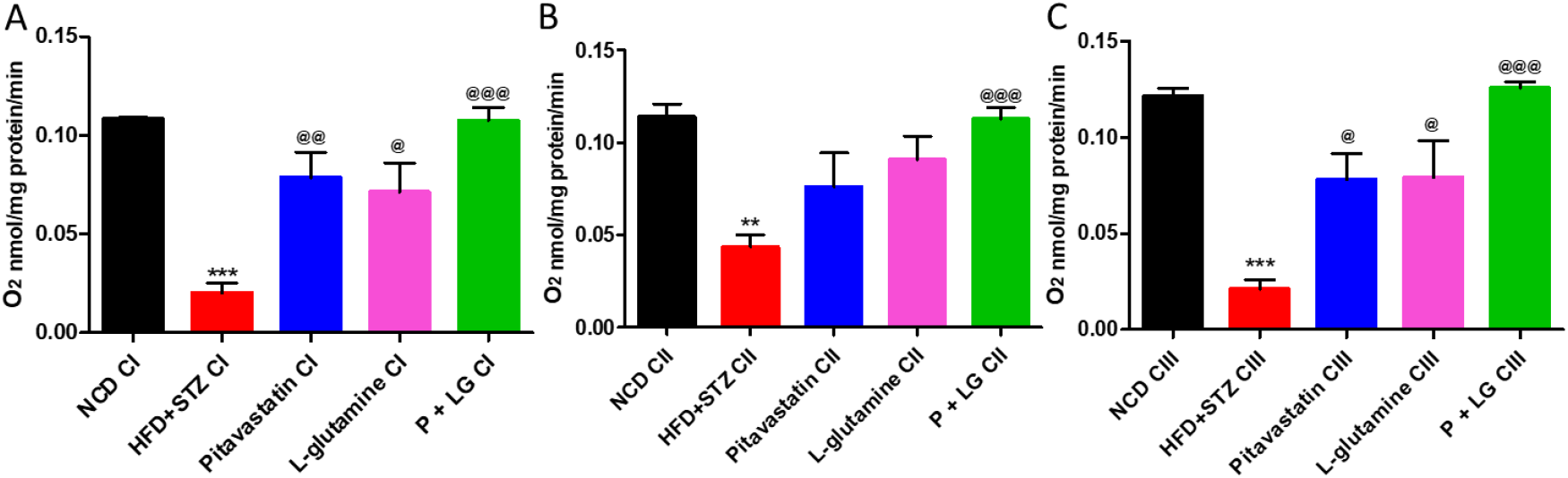
Ratio of oxygen consumption rate: **A) Complex I:** RCR of state 3/state 4 in HFD+STZ group was reduced as compared to NCD. There is a rise in the RCR in the P+LG treated group; **B) Complex II:** RCR of state 3/state 4 in HFD+STZ group is reduced as compared to NCD. There is a rise in the RCR in the P+LG treated group, and **C) Complex III:** RCR of state 3/state 4 in HFD+STZ group is reduced as compared to NCD. There is a rise in the RCR in the P+LG treated group. (^@^*p*<0.05, ^@@^*p*<0.01, ^@@@^*p*<0.001, vs. HFD+STZ, \*\**p*<0.01, \*\*\**p*<0.001, vs. NCD, n=3/group)

### 3.6 Western blot analysis

The levels of key insulin signaling pathway molecules were estimated by western blot. There was a significant decrease in the IR1β, pAKT/tAkt and AdipoR1 levels in the HFD+STZ treated group (^**^*p*<0.01, ^*^*p*<0.05 and ^**^*p*<0.01, respectively; Fig 8 B, C & E). The protein levels were resolved in the combination treated group (^@@@^*p*<0.001, ^@@^*p*<0.01 and ^@@^*p*<0.01, respectively; Fig 8 B, C & E). Further, there was a significant increase in the IR1β levels in pitavastatin and L-glutamine alone treated groups as compared to HFD+STZ (^@^*p*<0.05 and ^@@^*p*<0.01, respectively; Fig 8 B). However, no significant difference was observed in the pIRS/IRS ratio, GLUT 4 and PPARα levels (ns=non-significant; Fig 8 A, D and F).

**Figure 8:**
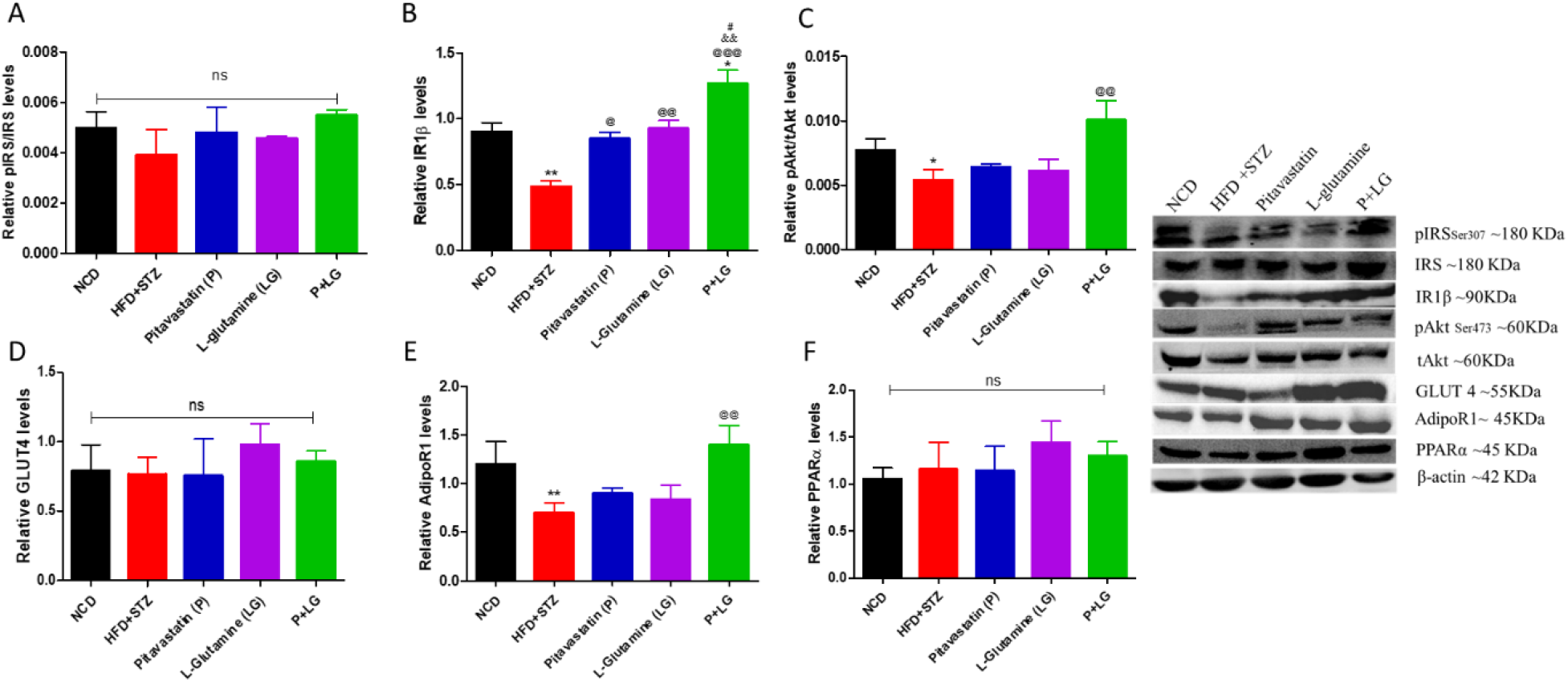
Insulin signaling pathway: **A) pIRS/IRS ratio:** There was no significant difference in the pIRS/IRS ratio amongst the various treated groups; **B) Insulin receptor1β:** There was a significant increase in the expression of IR1β in the combination treated group as compared to HFD+STZ group and individual treated groups; **C) pAKT/tAkt ratio:** There was a significant rise in the pAKT/tAkt ratio in the combination treated group as compared to HFD+STZ treated group**; D) GLUT4:** here was no significant difference in the GLUT4 levels amongst the various treated groups; **E) AdipoR1:** There was a significant rise in the AdipoR1 levels in the combination treated group as compared to HFD+STZ treated group; **E) PPARα:** There was no significant difference in the PPARα levels amongst the various treated groups. (^@@^*p*<0.01, ^@@@^*p*<0.001, vs. HFD+STZ, ^&&^*p*<0.01, vs. P, ^#^*p*<0.05, vs. LG, **\****p*<0.05, **\*\****p*<0.01, vs. NCD, n=3/group).

### 3.7 Pancreatic β-cells regeneration and apoptosis analysis

A significant decrease in the insulin positive β-cells and number of islets (data not shown) was observed in the HFD+STZ treated animals (^##^*p*<0.01; Fig 9 A panel, A’ & F). Upon treatment there was a significant rise in the Insulin/BrdU co-positive cells in Pitavastatin, L-glutamine and P+LG treated groups as compared to HFD+STZ (^**^*p*<0.01, ^*^*p*<0.05 and ^***^*p*<0.001, respectively; Fig 9 A panel and A’). Further, markers for neogenesis (PDX1 and NGN3), and transdifferentiation (PAX4 and ARX) were costained with insulin and it was observed that only PDX1 and PAX4 were costained with insulin (^**^*p*<0.01, ^***^*p*<0.001 & ^***^*p*<0.00, and ^*^*p*<0.05, ^***^*p*<0.001 and ^***^*p*<0.001, respectively; Fig 9 B panel & B’, and C panel & C’, respectively) upon mono- and combination treatment. No NGN3 and ARX positive cells were found. To check for apoptosis insulin/TUNEL and insulin/AIF staining was done. There was a significant increase in the insulin/TUNEL positive cells in the HFD+STZ treated group which was reversed upon the treatment with mono- and combination therapies (^***^*p*<0.001, ^***^*p*<0.001 & ^***^*p*<0.001; Fig 9 D panel and D’). However, no AIF translocation was observed in the HFD+STZ group (Fig 9 E panel). Overall, there was a significant increase in the number of Islets/pancreatic section in the combination treated groups as compared to HFD+STZ group (^***^*p*<0.001; Fig 9 F).

**Figure 9:**
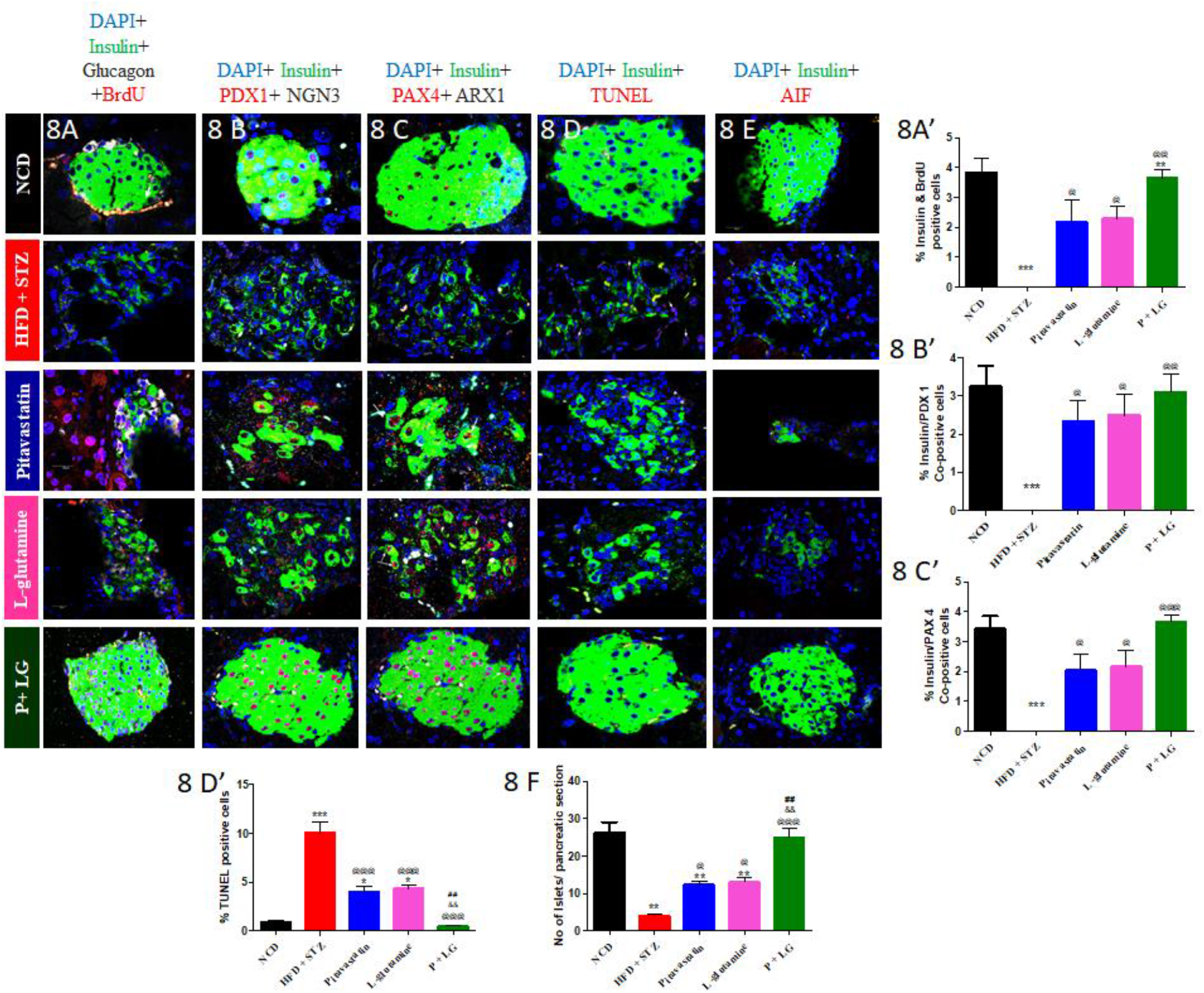
Immuno-histochemistry of pancreatic Islets of Langerhans at 60X: **A) Proliferation:** A significant increase in the Insulin/BrdU co-positive cells were observed in Pitavastatin, L-glutamine and P+LG treated groups as compared to HFD+STZ; **B) Neogenesis (Ins/PDX1/NGN3):** A significant increase in only Insulin/PDX1 co-positive cells was observed in Pitavastatin, L-glutamine and P+LG treated groups as compared to HFD+STZ; **C) Transdifferentiation (INS/PAX4/ARX1):** A significant increase in the Insulin/PAX4 co-positive cells were observed in Pitavastatin, L-glutamine and P+LG treated groups as compared to HFD+STZ; **D) Apoptosis:** A significant decrease in the Insulin/TUNEL co-positive cells were observed in Pitavastatin, L-glutamine and P+LG treated groups as compared to HFD+STZ; **E) AIF translocation:** No significant translocation of AIF was observed in any of the groups; **F) No. of islets/section:** A significant increase in no. of islets/section was observed in P+LG treated groups as compared to HFD+STZ, and AIF translocation was also not observed (^@^*p*<0.05, ^@@^*p*<0.01, ^@@@^*p*<0.001, vs. HFD+STZ, ^&&^*p*<0.01, vs. P, ^##^*p*<0.01, vs. LG,, \**p*< 0.05, \*\**p*< 0.01, \*\*\**p*< 0.001, vs. NCD, n=3/group, scale= 20µm, Magnification=60X)

## 4. Discussion

The data presented herein demonstrates that pitavastatin and L-glutamine in combination could ameliorate T2D by 1) improving glucose tolerance and insulin sensitivity, 2) normalizing insulin and adiponectin concentrations, 3) inhibiting gluconeogenesis and glycogenolysis, reducing hepatic GLUT2 and enhancing glycolysis, glycogen synthesis, 4) increasing IR1β, pAkt, AdipoR1 expression levels in skeletal muscle, 5) increasing mitochondrial biogenesis and its complex activities, and regenerating β-cells (Fig 10).

**Figure 10:**
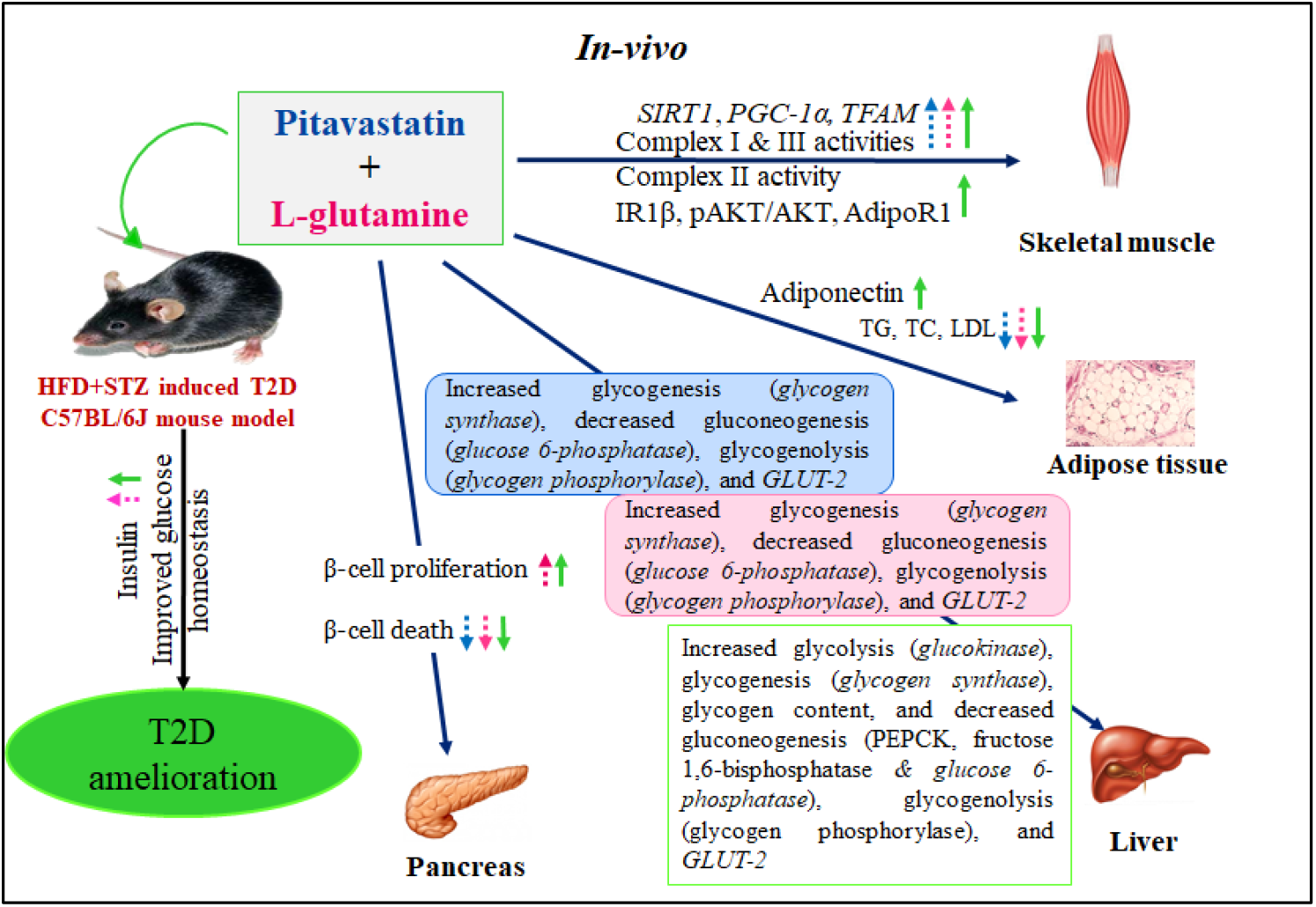
Pitavastatin and L-glutamine could induce normolipidemia and reduce the transcript levels of glucose 6-phosphatase, glycogen phosphorylase and GLUT2 in liver whilst increasing SIRT1, PGC1α and TFAM, mitochondrial complex I and III activities in skeletal muscle. Both pitavastatin and L-glutamine could also reduce β-cell death individually and L-glutamine also could induce β-cell proliferation. In combination they could additionally increase mitochondrial complex III activity in skeletal muscle, and glucokinase transcript levels with a concomitant decrease in PEPCK and fructose 1,6-bisphophatase activities in liver. The combination also induced normoadiponectinemia, increased phosphorylation of Akt, increase IR1β and AdipoR1 protein levels in skeletal muscle, and insulin levels. As a whole the drugs could work together synergistically to bring normolipidemia, reverse mitochondrial dysfunction, decrease gluconeogenesis, glycogenolysis, glycogen synthesis and GLUT 2. The combination therapy could also reduce β-cell death, increase insulin levels, enhance glucose tolerance and insulin sensitivity thus being able to ameliorate T2D in HFD+STZ induced mouse model.

Obesity has been strongly associated with the reduction of anti-inflammatory and increase of pro-inflammatory adipokines. Many of these adipokines are crucial in maintaining the status of insulin sensitivity of peripheral tissues viz. adipose tissue and skeletal muscle (Pramanik et al., 2017, Patel et al., 2016; Patel et al., 2018; Patel et al., 2019; Rathwa et al., 2019 a & b; Rathwa et al., 2020). In view of this, regulating hyperglycaemia by merely regulating the hepatic glucose out-put or enhancing glucose clearance seem superficial. Moreover, as previously stated, T2D also shows significant reduction in β-cell mass. Thus, addressing the white elephant in the room i.e., glucolipotoxicity and loss of β-cells in T2D we chose two drugs: pitavastatin and L-glutamine. We chose pitavastatin as it is unique in engaging pathways that not only lowers blood cholesterol levels, but also excess cellular cholesterol levels by virtue of its pharmacological profile and enhances adiponectin in hyperlipidaemic T2D conditions. In our HFD+STZ induced mice model, we found pitavastatin to be effective in bringing about glucose homeostasis, increasing adiponectin levels and reducing TG, TG and LDL as supported by previous studies reporting statins to enhance glucose tolerance and increase anti-inflammatory adipokines by reducing LDL and c-reactive proteins (Lee et al., 2016; Yoshika et al., 2010; Matsubara et al., 2012; Iwata et al., 2019; Chen et al., 2019; Cho et al., 2020). Evidence from *in vitro* studies suggest that the off-target effect of pitavastatin on adiponectin may be related to the prevention of adipocyte hypertrophy and adipokine dysregulation (Ishihara et al., 2010). L-glutamine treated group also showed enhanced glucose homeostasis (Greenfield et al., 2009; Molfino et al., 2009; Samocha-Bonet et al., 2011), increased adiponectin levels (Abboud et al., 2019) and reduced TG, TC and LDL levels (Alba-Loureiro et al., 2009; Badole et al., 2013). We also found increased insulin in the L-glutamine treated group as reported earlier (Abboud et al., 2019; Badole et al., 2013). Supporting our findings, liver in T2D model when infused with L-glutamine is reported to significantly increase L-alanine production, which is consumed at a high rate in β cells exerting insulinotropic effects (Dixon et al, 2003; Newsholme et al., 2006). In the combination treatment, we saw an additive effect of enhanced insulin levels, normoadiponectinemia, glucose tolerance, insulin sensitivity, FBG levels and improved lipid profile. Gluco- and lipotoxicity is known to induce stress and mitochondrial dysfunction. In addition, statins have been associated with impaired mitochondria, decreased oxidative phosphorylation capacity, decreased mitochondrial membrane potential and activation of intrinsic apoptotic pathway in many studies (Galtier et al., 2012; Sirvent et al., 2012). On the contrary, glutamine has been reported to support respiration in an *in-vitro* study where it was reported to reduce cell permeability and cytochrome c levels, and increase mitochondrial membrane potential, maintaining mitochondrial integrity (Ahmad et al., 2001; Safi et al., 2015). Hence, we studied the transcript levels of key mitochondrial biogenesis markers and the mitochondrial complex (I, II & III) activities. Pitavastatin could increase the transcript levels of *PGC1α, TFAM* and *SIRT1*, thus alleviating oxidative stress, and enhance the electron transport chain (ETC) complexes I and III activities as reinforced by previous studies (Ota et al., 2010, Vevera et al., 2016). Our results for the first time suggest that L-glutamine can also induce mitochondrial biogenesis and increase complex (I and III) activities. This is probably an effect of reduced oxidative stress as induced by L-glutamine, a potent antioxidant (Tsai et al., 2012; Badole et al., 2013; Badole et al., 2014). We saw an additive effect of P+LG treatment on mitochondrial biogenesis and its complex (I, II & III) activities. Moreover, the glucoregulatory pathway was also significantly corrected upon pitavastatin treatment with an increased expression of *GCK* mRNA levels and its activity, increased glycogenesis and glycogen content supported by Daido et al., 2014. Further, we found decreased insulin resistance and hepatic *GLUT2* mRNA expression. Supporting our results Fraulob et al. too had reported HFD fed C57BL/6 mice treated with rosuvastatin ameliorated insulin resistance and hepatic steatosis. Moreover, a down regulation of hepatic *GLUT-2* expression was observed in statin-treated mice (Fraulob et al., 2012) suggesting statin may not increase glucose overproduction in already hyperlipidemic mouse. In addition, we also found a concomitant decrease in the PEPCK and FBPase mRNA expression levels and activities. The fact that pitavastatin increased adiponectin levels makes it more conspicuous as to how it could mediate insulin sensitivity. Adiponectin is known to suppress hepatic glucose output and lower systemic glucose by enhancing hepatic insulin sensitivity, and inhibiting expression and activity of gluconeogenesis key enzymes (Turer et al., 2012; Miller et al., 2011). L-glutamine was also found to enhance the glucoregulatory pathway by reducing hepatic glucose output (Abboud et al., 2019). The cleaved GLP-1 product (9–36 amino acids), has been reported to suppress gluconeogenesis and act as an antioxidant (Thomas and Habener, 2010). Further, the administration of L-glutamine could reverse the defective glucoregulatory pathway by increasing glycogen synthesis as reported previously (Varnier et al., 1995) and reducing *GLUT2* mRNA expression as *glycogen phosphorylase* mRNA expression was reduced. Bringing the monotherapies together in combination therapy, it could enhance the hepatic glucoregulatory pathway by inhibiting gluconeogenesis, reducing GLUT2 expression, and increasing glycolysis. Previous studies with atorva-, prava-, simva- or rosuvastatin showed the suppression of the phosphorylation of IR, IRS-1 and Akt, and total expression of IR, IRS-1 or glycogen synthase kinase-3β (GSK-3β), but not Akt, in L6 skeletal muscle myotubes and C2C12 myotubes (Li et al., 2016; Yaluri et al., 2016). In this context we further went on to study the effect on insulin signalling pathway and found a significantly enhanced IRS-1/phosphatidylinositol 3-kinase/Akt pathway leading to glucose uptake in mono- and additively in combination therapies. We hypothesize these effects to be a downstream effect of the increased adiponectin levels (Ishihara et al., 2010). This is the first report wherein the effect of pitavastatin on insulin signalling pathway was studied *in-vivo*. Going hand in hand with our data, in the latest in-vitro study it was found that there was an increased insulin stimulated GLUT4 translocation onto the membrane of 3T3-L1 cells on pitavastatin treatment (Cho et al., 2020). Similarly, in Wistar rats fed with HFD with/without lovastatin showed enhanced insulin-stimulated IRS-1/phosphatidylinositol 3-kinase/Akt pathway in skeletal muscle from lovastatin-treated rats (Lalli et al., 2008). Prior reports on L-glutamine also suggest its insulin signalling enhancing properties (Wang et al., 2018; Abboud et al., 2019). Finally, our immunohistochemistry data suggests a significant reduction in β-cell death in both mono- and combination therapies. Further, pitavastatin and L-glutamine could induce proliferation as the insulin positive β-cells expressed PDX1 and PAX4 along with a significant incorporation of BrdU in the nucleus. In a recent *in-vitro* experiment it was found that most β-cells treated with low dose of pitavastatin were in S and G_2_/M phase as compared to other statin groups (Zhao and Zhao, 2015). L-glutamine controls the biosynthesis of IGF2 which is an autocrine regulator of β-cell mass and function. It also activates Akt phosphorylation in β-cells leading to its proliferation (Moullé et al., 2017). L-glutamine has also been reported to be able to upregulate PDX1 (Corless et al., 2006). PDX1in concert with NGN3 and MAFA is an important marker of islet neogenesis (differentiation of acinar cells to β-cell cells) while PAX4 is a marker of transdifferentiation (α-cell to β-cell conversion) (Zhu et al., 2015; Zhang et al., 2016). However, we did not find NGN3 and insulin co-positive cells. Hence, the data here indicates a significant rise in β-cells via proliferative pathway alone. Though there was only a marginally significant increase in the number of islets in pitavastatin and L-glutamine treated groups, they showed an increase in PDX1 and PAX4 insulin co-positive cells demonstrating their individual β-cell regenerative properties. PAX4 was also reported to promote differentiation of PDX1 positive mesenchymal stem cells to β-cell fate (Xu et al., 2017). Future studies in this direction are warranted. Three large meta-analyses of randomized, controlled trials showed those who are >60 years of age or are being treated with an intensive-dose of statin therapy have higher chances of developing new-our results (Rajpathak et al., 2009; Sattar et al., 2010; Preiss et al., 2011) clearly stating that effect of statin is largely based on the dosage recommended. Though several preclinical and clinical studies have been carried out till date to explore the individual therapeutic potential of pitavastatin and L-glutamine to ameliorate T2D this is a first attempt to study the combination effect of the two drugs on various diabetes manifestations. A point to worth mentioning here is that combination of DPP4 inhibitor sitagliptin (extends GLP-1 half-life) in combination with simvastatin was approved by the US Food and Drug Administration in October 2011 and sold under the brand name Juvisync. It is a fixed-dose combination anti-diabetic medication used to treat T2D and hypercholesterolemia (Ramadan et al., 2015).

## 6. Conclusion

Pitavastatin and L-glutamine in combination could ameliorate T2D acting additively at hepatic, skeletal muscle and pancreatic fronts.

## Abbreviations

T2D: Type 2 Diabetes
FBG: Fasting Blood Glucose
TC: Total Cholesterol
HDL: High Density Lipoprotein
TG: Triglycerides
LDL: Low Density Lipoprotein
BMI: Body Mass Index
HFD: High Fat Diet
STZ: Streptozotocin
LG: L-glutamine
PDX1: Pancreatic and Duodenal Homeobox 1
ARX1: Aristaless-related homeobox-encoding gene
PAX4: Paired box 4
NGN3: Neurogenin 3

## 7. Acknowledgments

We thank Dr. Deepak Sharma, BARC, Mumbai for providing the confocal facilities. NR thanks University Grants Commission-National Fellowship for higher education for ST students, New Delhi, India, for awarding SRF. RP thanks Council for Scientific and Industrial Research, New Delhi, India for awarding SRF.

## 10. Author Contributions

RB conceived the idea and designed the experiments. SPP performed the experiments, data acquisition, data analysis and wrote the manuscript; RP, NP and NR helped in data acquisition and data analysis. RB and AVR contributed to the critical revision of the manuscript.

